# The Quorum Sensing Peptide EntF* Promotes Colorectal Cancer Metastasis in Mice: A New Factor in the Microbiome-Host Interaction

**DOI:** 10.1101/2020.09.17.301044

**Authors:** Nathan Debunne, Evelien Wynendaele, Yorick Janssens, Anton De Spiegeleer, Frederick Verbeke, Liesa Tack, Sophie Van Welden, Evy Goossens, Daniel Knappe, Ralf Hoffmann, Christophe Van De Wiele, Debby Laukens, Peter Van Eenoo, Filip Van Immerseel, Olivier De Wever, Bart De Spiegeleer

**Author notes:** These authors contributed equally: Nathan Debunne, Evelien Wynendaele. **Corresponding author**, Bart De Spiegeleer, Drug Quality and Registration Group, Faculty of Pharmaceutical Sciences, Ghent University, Ottergemsesteenweg 460, B-9000 Ghent, Belgium.

## Abstract

**Background:** Colorectal cancer, one of the most common malignancies worldwide, is associated with a high mortality rate, mainly caused by metastasis. Comparative metagenome-wide analyses between healthy individuals and cancer patients suggest a role for the human intestinal microbiota. Nevertheless, which microbial molecules are involved in this communication is largely unknown, with current studies mainly focusing on short chain fatty acids and amino acid metabolites as potential mediators. However, quorum sensing peptides are not yet considered in this microbiome-host interaction: their *in vivo* presence nor any *in vivo* host-effect have been reported.

**Results:** For the first time, we showed that a quorum sensing peptide metabolite, EntF* produced by intestinal microbiota (*E. faecium*), is present in the blood circulation of mice. Moreover, it significantly promotes colorectal cancer metastasis *in vivo*, with metastatic lesions found in both liver and lung tissues, using an orthotopic mice model evaluating bioluminescence as well as macroscopic and microscopic presence of metastatic tumour nodules. *In vitro* tests on E-cadherin expression levels thereby indicated that the first, second, sixth and tenth amino acid of EntF* were critical for the epithelial-mesenchymal transition (EMT) effect, responsible for tumour metastasis.

**Conclusion:** This paper adds a new group of molecules, the quorum sensing peptides, as an additional causative factor explaining the microbiome-host interaction. The presence of a selected quorum sensing peptide (metabolite) in the mouse was proven for the first time and its *in vivo* effect on colorectal metastasis was demonstrated. We anticipate our *in vivo* results to be a starting point for broader microbiome-health investigations, not only limited to colorectal cancer metastasis, but also for developing novel bio-therapeutics in other disease areas, giving due attention to the QSP produced by the microbiome.

## BACKGROUND

Colorectal cancer (CRC) is the third most common malignancy worldwide and associated with a high mortality rate, mainly caused by metastasis (mCRC). Primary CRC originates from epithelial cells that line the gastrointestinal tract, usually (but not always) through an adenoma-carcinoma sequence in the CRC tumorigenesis^1^: normal colorectal epithelium transforms to an adenoma and ultimately to an invasive and metastatic tumour. In a first step of the metastasis process, the epithelial CRC cells switch towards a mesenchymal phenotype, known as epithelial-to-mesenchymal transition (EMT)^2^. Although a clear hereditary component in CRC tumorigenesis is present in some cases, a strong association with diet and lifestyle has been demonstrated as well^3^. Moreover, also inflammation is believed to play a role in CRC cancer development, as a driver, illustrated by colitis-associated colon (CAC) cancers in patients with inflammatory bowel diseases (IBD), as well as a consequence, seen in patients with sporadic colorectal cancer^4^.

Growing evidence obtained in the last decade also suggests a role for the human intestinal microbiota in CRC^5–7^. For example, by comparing faeces from healthy persons and patients, higher abundances of *Enterococcus, Escherichia* and *Fusobacterium* species were observed in multiple intestinal disorders, including colorectal cancer (CRC) and Crohn’s disease^8–11^. However, the causative factors for disease development or progression are not well understood and current research is mainly limited to bacterial-derived short-chain fatty acids and amino acid-derived amines^12^.

Quorum sensing peptides are traditionally regarded as intra- and inter-bacterial communication molecules; however, given their wide structural variety and co-evolution, we anticipate that these bacterial metabolites may also interact with the host. Different quorum sensing peptides were indeed previously found to influence the behaviour of the host cells, going from cancer cells (colorectal and breast cancer) towards brain and muscle cells^13–16^. In colorectal cancer, specific microbial quorum sensing peptides were found to promote tumour cell invasion and angiogenesis *in vitro*, indicating the possible pro-metastatic properties of these peptides. In this study, we focus on *Enterococcus faecium*, one of the most abundant species in the human intestinal microbiota, which synthesizes the enterocin induction factor, *i.e*. the propeptide of the EntF quorum sensing peptide (AGTKPQGKPASNLVECVFSLFKKCN). This peptide serves as a communication signal, regulating the production of enterocin A and B toxins, which are produced to inhibit the growth of similar or closely related bacterial strains^17–22^.

Up till now, however, quorum sensing peptides have not yet been unambiguously demonstrated to be present in biofluids. Only an indirect indication of the *in vivo* presence of an unidentified quorum sensing peptide was described in the stool of patients suffering from a *Clostridium difficile* infection^23^. Indicating the biological presence of certain quorum sensing peptides in mice, together with their *in vivo* effect on the host, may stimulate the research towards the additional role of quorum sensing peptides in the microbiome-host interaction.

## RESULTS AND DISCUSSION

### Biological presence of EntF* in mice

*In vitro* metabolization studies of EntF (Fig. 1a) in faeces and colonic mice tissue homogenates quickly yielded a 15-mer peptide EntF* (SNLVECVFSLFKKCN) (Fig. 1b), with a mean (± s.e.m.) formation rate of 1.71 (± 0.27)% min^−1^ and 0.11 (± 0.01)% min^−1^, respectively. Similar to other quorum sensing peptides^14^, this EntF* peptide is also able to cross the intestinal barrier *in vitro*, using a CaCo-2 monolayer permeability assay, with a mean (± s.e.m.) apparent permeability coefficient of 3.70 (± 0.22) x 10^−9^ cm s^−1^ (Fig. 1c). These *in vitro* studies thus indicated that EntF* can be present in the blood circulation of the host, *i.e*. after degradation of the endogenously present EntF to EntF* in the colon or faeces and subsequent intestinal absorption of the EntF* peptide.

**Figure 1:**
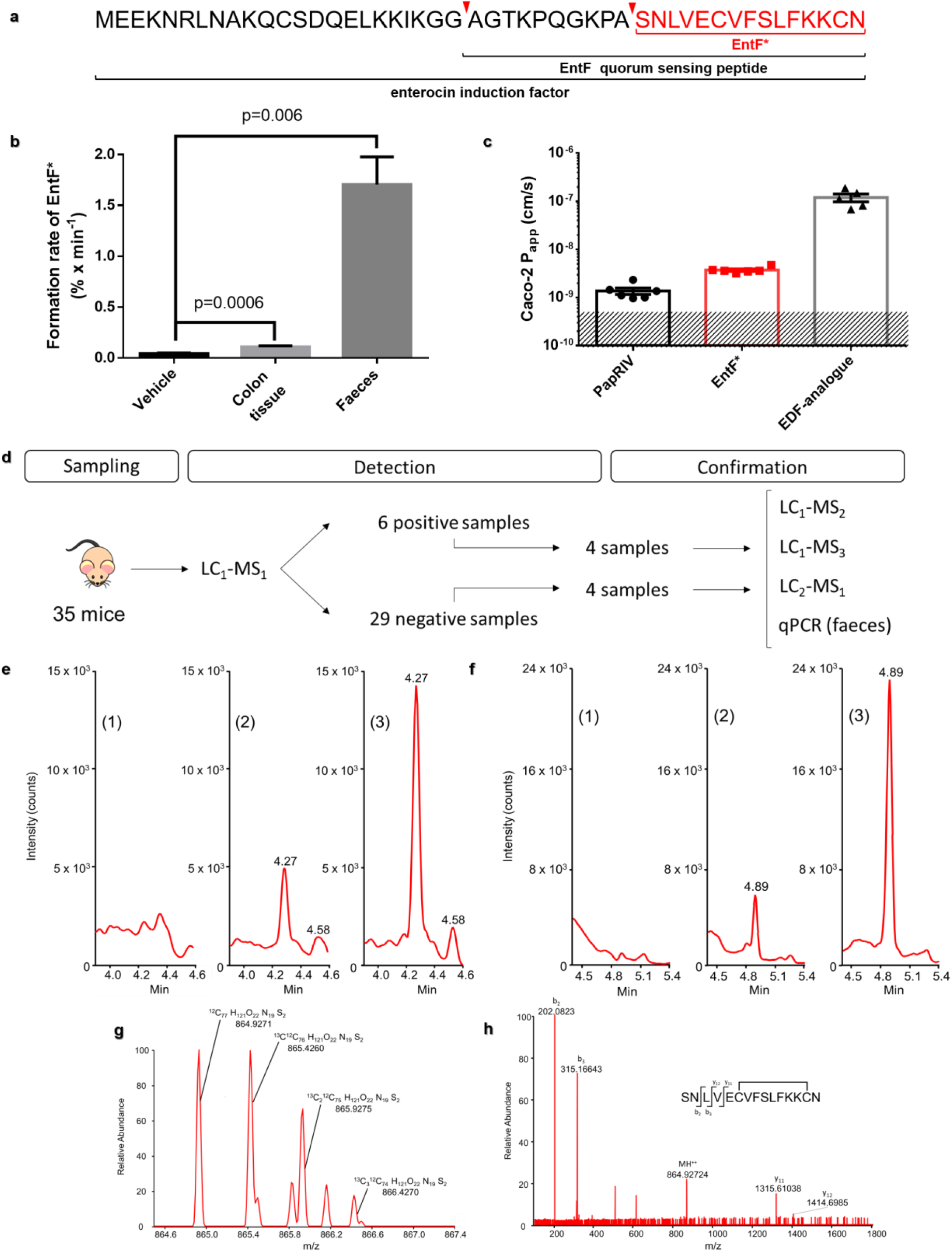
*In vitro* formation and *in vivo* presence of the EntF quorum sensing peptide-derived metabolite. **a,** Sequence of the quorum sensing propeptide enterocin induction factor, the mature quorum sensing peptide EntF and its metabolite EntF*. **b,** The *in vitro* formation rate of EntF* from EntF in colon and faeces homogenate. Bars represent mean formation rate ± s.e.m from 6 (colon), resp. 4 (faeces) independent experiments. Statistically significant differences were determined by a Mann-Whitney U test with indicated p-values. **c,** CaCo-2 apparent permeability coefficients (P_app_) of 3 different quorum sensing peptides. Bars represent mean P_app_-values ± s.e.m. (n=5-6 independent experiments per group); the shaded area represents the limit of detection. **d,** Flow chart of *in vivo* data acquisition, from sampling of serum samples to detection and confirmation of EntF*. Different LC-MS methods: reversed-phase ultra-high-performance liquid chromatography (RP-UPLC) using triple quadrupole (TQ) in MRM mode (LC_1_-MS_1_), high-resolution quadrupole time-of-flight (LC_1_-MS_2_), high-resolution quadrupole-orbitrap (LC_1_-MS_3_) and HILIC-amide UPLC using TQ in MRM mode (LC_2_-MS_1_). qPCR was performed on faeces samples from those mice to demonstrate the presence of EntF*-containing *E. feacium* DNA copies. **e,** Chromatographic profile of (1) negative serum sample; (2) positive serum sample; (3) serum of mice i.p. injected with EntF*; all using RP-UPLC with detection by electrospray ionization mass spectrometry (ESI-MS) using TQ in MRM mode (*m/z*= 865 → 202.08 + 315.17). **f,** Chromatographic profile of (1) negative serum sample; (2) positive serum sample; (3) serum of mice i.p. injected with EntF*; all using HILIC amide UPLC with detection by ESI-MS using TQ in MRM mode (*m/z*= 865 → 202.08 + 315.17). **g,** Isotopic distribution of the double charged EntF* measured in a positive serum sample using RP-UPLC with detection by ESI-MS using quadrupole-orbitrap. **h,** High-resolution tandem mass spectrum of EntF* with characteristic fragments, using RP-UPLC with detection by Q-TOF.

To unambiguously demonstrate the *in vivo* presence of EntF*, a bioanalytical method using reversed phase ultra-high-performance liquid chromatography coupled to a triple quadrupole mass spectrometer (RP-UPLC-TQ-MS) in Multiple Reaction Monitoring (MRM)- mode was developed and optimized, aiming to avoid carry-over and adsorption, as well as to maximize the selectivity and sensitivity. Critical methodological aspects to achieve these goals were: (1) a suitable sample preparation method using a novel bovine serum albumin (BSA)- based anti-adsorption solution^24^ and the combination of solvent/acid/heat sample treatment followed by solid phase extraction, and (2) appropriate MS detection settings, including the selection of quantifier (b_2_: *m/z*= 202.08) and qualifier (b_3_: *m/z*= 315.17) ions. The method was suitably verified and found to be appropriate for its purpose (Supplementary Fig. 1). Serum samples of 35 healthy, non-manipulated mice (C57BL/6 mice, aged 5-18 months) were collected and analysed for the endogenous, natural presence of EntF* (Fig. 1d), using the developed RP-UPLC-TQ-MS (LC_1_-MS_1_) method. For six mice, the presence of EntF* was observed in their serum above the limit of quantification (LOQ) of 100 pM (Fig. 1e). Taking into account all mice results, *i.e*. including the <LOQ (zero) values, an overall estimated mean value of 305 pM (s.e.m.=138 pM; n=35) was obtained (Supplementary Table 1). Following these findings, further evidence was obtained by subjecting a selected set of samples to three additional chromatographic methods: Hydrophilic Interaction Liquid Chromatography (HILIC)-UPLC-TQ-MS (LC_2_-MS_1_) (Fig. 1f) as an orthogonal separation system, and RP-UPLC-QTOF-MS (LC_1_-MS_2_) and RP-UPLC-QOrbitrap-MS (LC_1_-MS_3_) as high-resolution mass spectrometers. Serum samples of eight mice (four positive and four negative samples (*i.e*. above and below the LOQ, respectively, based on the RP-UPLC-TQ-MS findings)) were analysed using these additional methods (Fig. 1f-h). The presence and identity of EntF* was confirmed using the isotopic distribution of the doubly charged precursor ion (Fig. 1g) and the presence of fragment ions y_11_ (*m/z*= 1315.61) and y_12_ (*m/z*= 1414.69) (Fig. 1h) in the four positive serum samples (Supplementary Table 1). Finally, quantitative real-time PCR analysis on the associated faeces samples was performed to demonstrate the existence of EntF*-containing *E. faecium* DNA copies (Supplementary Fig. 2, Supplementary Table 1): EntF* DNA copies were indeed observed in all four positive samples. The faeces samples that contained the EntF* gene but tested negative during the serum UPLC-MS analyses (*e.g*. sample 20181011S8) could possess specific *E. faecium* strains that show a reduced translational efficiency. This is also observed in the data that are presented in Supplementary Table 2: out of the three *E. faecium* strains that contain the EntF gene, only one strain produced EntF *in vitro* (LoD = 1.5 nM). In addition, standard protein BLAST searches indicated no endogenous presence of the EntF* peptide sequence in the murine genome (maximum sequence alignment of 67%), which is again a strong indication of the microbial origin of the *in vivo* found EntF* peptide.

### In vitro activity and molecular target of EntF*

EntF* was previously found by our group to selectively promote angiogenesis and tumour cell invasion in screening *in vitro* experiments using HCT-8 colorectal cancer cells^13,14^. These *in vitro* effects were now confirmed and extended. Using Western blotting, EntF* and some alanine- or D-amino acid derived analogues affected E-cadherin expression, which is linked to the epithelial-mesenchymal transition (EMT) of cancer cells (Fig. 2a, 2b and 2c, Supplementary Fig. 3). A mean significant decrease of 38% in E-cadherin expression was determined for EntF*. When the first, second or tenth amino acid of EntF* was replaced by an alanine amino acid, this decrease was significantly flattened out. These three amino acids are thus important for their contribution to the EMT-promoting effects of EntF*, while the other amino acids contribute to a much lesser amount, as determined by the Fisher’s LSD p-values and confirmed using the Jenks natural break algorithm. Replacing the sixth amino acid of EntF* by its unnatural D-amino acid isomer did also restore the E-cadherin expression to placebo levels. These results thus indicate that also the stereochemical configuration of the sixth amino acid of EntF* is of importance for its EMT-promoting effects. Moreover, by using the antagonist Nef-M1 for the CXCR4 receptor, together with EntF* on HCT-8 colorectal cells, the E-cadherin expression level increased from 62% to 94% (Cohen’s d effect size of 1.2), indicating the interaction of EntF* with the CXCL12 (or SDF-1)/CXCR4 pathway in tumour metastasis (*i.e*. EMT promotion). Interestingly, the modified peptide EntF* 1A, where the serine amino acid at position 1 of EntF* is replaced by an alanine amino acid, could also be identified as having an antagonistic activity towards EntF* on the CXCR4 receptor: E-cadherin expression levels increased from 62% to 92% (Cohen’s d effect size of 0.8) when EntF*1A was added to the EntF*-treated cells (Fig. 2d).

**Figure 2:**
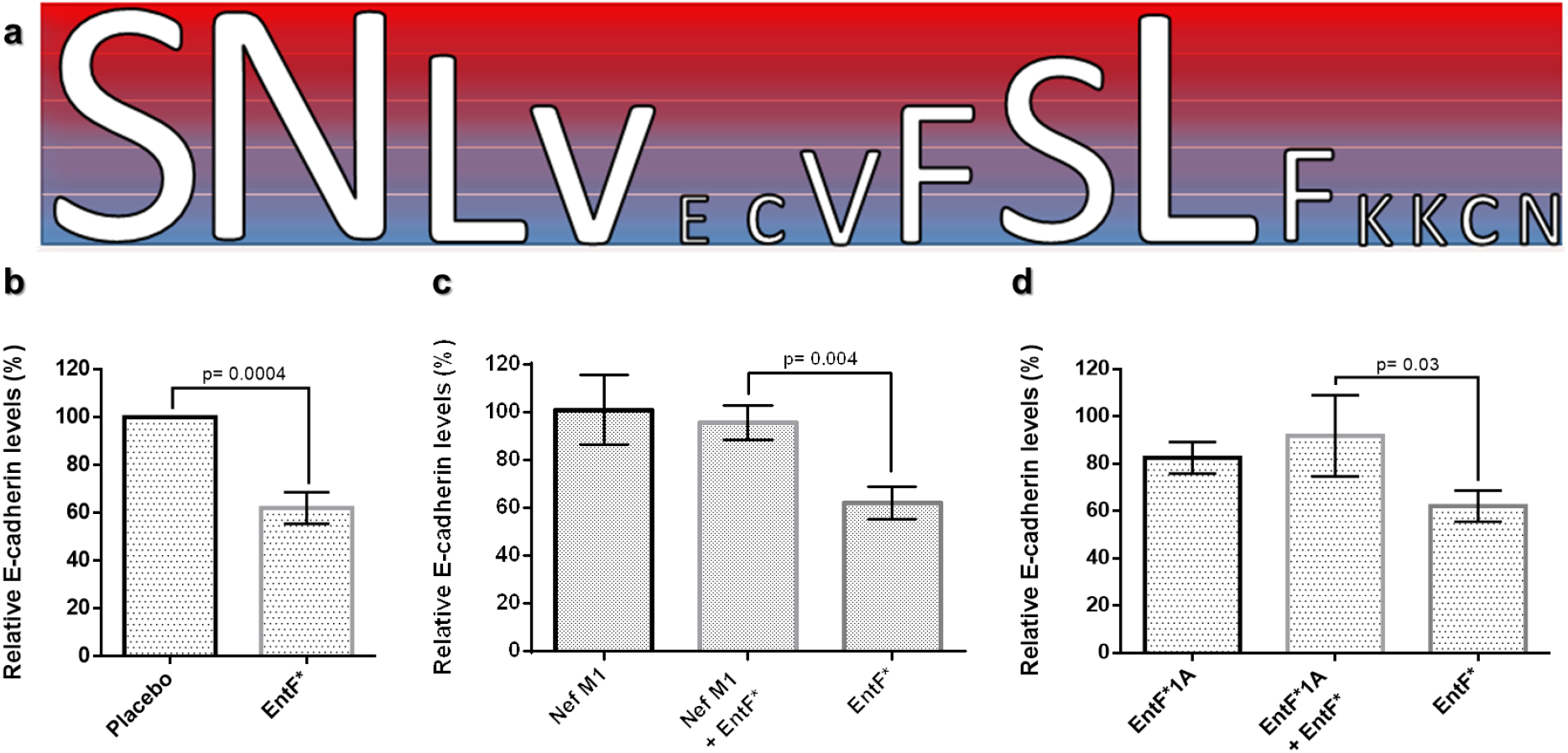
*In vitro* activity of the EntF* peptide. **a,** Effect of alanine-derived EntF* analogues on E-cadherin expression. Ranking in five classes (blue to red: increasing significance) was performed using the Fisher’s LSD p-values, which was confirmed using the Jenks natural break algorithm. Based on ranking, it was proven that the first, second and tenth amino acid of EntF* are the most important amino acids for the epithelial-mesenchymal promoting (EMT) effects of EntF*. **b,** Effect of EntF* on E-cadherin expression. A significant mean decrease of 38% in E-cadherin level for EntF* in comparison with placebo was observed (one-way ANOVA, Fisher’s LSD). **c,** The antagonistic effects of Nef-M1 on the E-cadherin reducing effect of EntF* on HCT-8 cells. Statistically significant differences were determined by a one-sided student’s t test. **d,** The antagonistic effects of EntF*1A on the E-cadherin reducing effect of EntF* on HCT-8 cells. Statistically significant differences were determined by a one-sided student’s t test.

### In vivo pro-metastatic properties of EntF* in mice

Having demonstrated the presence of EntF* *in vivo* as well as its *in vitro* effects, we evaluated its *in vivo* metastasis-promoting activities using an orthotopic colorectal cancer mouse model^25,26^. Before the luciferase-transfected HCT-8 cells were implanted into the wall of the caecum of the mice, cells were treated daily for five days with EntF* (100 nM), phosphate-buffered saline (PBS) vehicle or Transforming Growth Factor α (TGFα, positive control) (0.1 μg mL^−1^). On the sixth day, 6-weeks-old female Swiss nu/nu mice were orthotopically injected with the luciferase-transfected colorectal cancer cells, followed by a once-daily i.p. treatment of EntF* (100 nmol kg^−1^), PBS vehicle or Epidermal Growth Factor (EGF, positive control) (100 μg kg^−1^) (Fig. 3a). The *in vivo* distribution profile of EntF* in these mice was then determined, after which the EntF* daily exposure after i.p. treatment of 100 nmol kg^−1^ was calculated. Additionally, the natural endogenous daily exposure was calculated from the obtained endogenous EntF* levels, described in supplementary Table 1. Based on both exposure calculations, it could be concluded that daily injections of 100 nmol kg^−1^ EntF* gave daily peptide exposures which were five times higher than the endogenous (natural) exposure in those mice, hence, in the range of the “positive” mice, thus demonstrating the biological relevance of this experimental set-up (Fig. 3b). Bioluminescent imaging of the mice was performed weekly to monitor the tumour growth (Fig. 3c). During the course of 6 weeks, EntF* caused a statistically significant increase in luciferase activity compared to vehicle (p=0.030). This increase was even not significantly different from the well-established positive control EGF (p=0.319; Fig. 3d). Our results thereby demonstrated, after 6 weeks treatment, an effect size of 128% increase in bioluminescence for EntF* compared to the placebo PBS, varying from −75%, a relative small negative association, to a 1494% increase, a substantial positive association (Fig. 4a). For the positive control EGF, a median effect size of 316% was obtained, ranging from −145% to 850%. This was confirmed by the number of tumour nodules counted macroscopically on the caecum, which was again statistically significantly higher with EntF* in comparison to PBS (p=0.036) (Fig. 3d-e), demonstrating the *in vivo* 3-fold increase in the number of nodules due to this quorum sensing peptide metabolite while EGF showed a 4.5-fold increase (Fig. 4b).

**Figure 3:**
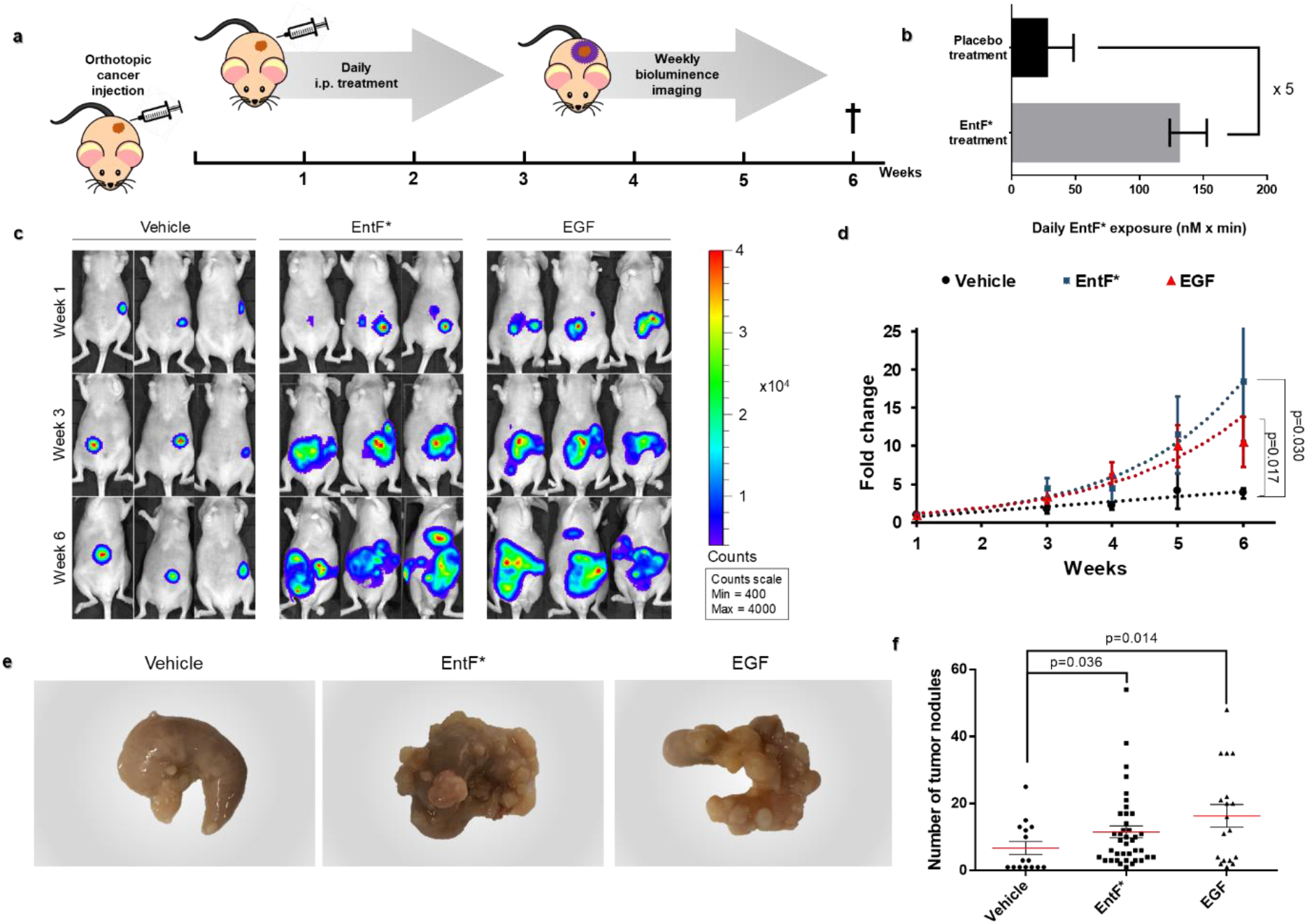
*In vivo* metastasis-inducing effect of EntF* in an orthotopic colorectal cancer mouse model. **a,** Experimental schematic timeline. Female Swiss nu/nu mice were orthotopically injected with 1 x 10^6^ luciferase transfected HCT-8 cells at the age of 5 weeks. During 6 weeks, the mice were daily i.p. injected with 100 nmol kg^−1^ EntF*, PBS control or 0.1 mg kg^−1^ EGF positive control. Bioluminescent imaging was performed weekly to determine cancer progression. After 6 weeks, the mice were euthanized and the caecum, liver and lungs collected. **b,** The daily exposure in the female Swiss nu/nu mice (n=65 placebo mice, with negative mice (< LoQ = 100 pM) set as 0 pM) (black) and the female Swiss nu/nu mice after injection with EntF* (gray, n=14) is given. Error bars represent s.e.m. values. It could be concluded that the daily exposure after i.p. injection of 100 nmol kg^−1^ EntF* (used for the orthotopic colorectal cancer mouse model) is 5 times higher than the natural occurring EntF* levels. **c,** A representative image comparing the basal bioluminescence activity between the three treatments. Mice were i.p. injected with 150 mg kg-1 luciferine and imaged 10 minutes later in the supine position. **d,** Tumour growth curves of the three groups. Based on linear regression slope comparison, the EntF* as well as the positive control EGF treatment resulted in a significant increase of tumour growth compared to the vehicle control with indicated p-values. Data represent mean fold change ± s.e.m. (n= 17-18 mouse per group). **e,** Macroscopic, representative caecum pictures of the three treatments at the end of the experiment. **f,** Caecum tumour nodules were counted and the data represent the mean ± s.e.m. Statistically significant differences were determined by a Mann-Whitney U test (n= 15-38 per group) with indicated p-values.

**Figure 4:**
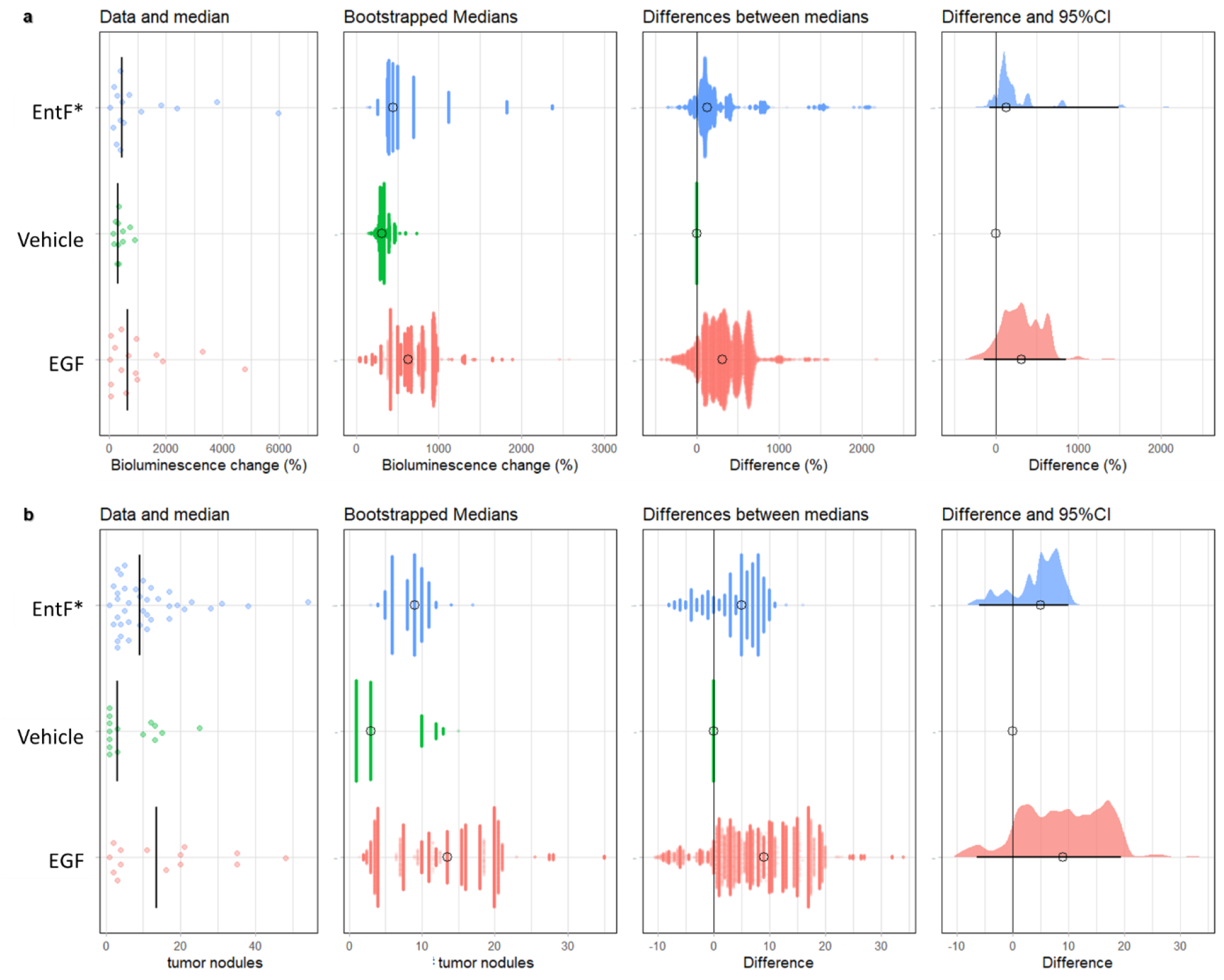
Effect size for bioluminescence and the number of nodules on the caecum in the orthotopic mouse model after 6 weeks treatment. **a,** After 6 weeks treatment, an effect size of 128% increase in bioluminescence for EntF* compared to the placebo PBS was observed, while for the positive control EGF, a median effect size of 316% was obtained. When calculating the effect size according to Hedges’ G values, a median to high effect was observed for both the EntF* and the EGF treatment groups, compared to the placebo group. **b,** After 6 weeks treatment, a 3-fold increase in the number of nodules on the caecum was observed for EntF* compared to the placebo PBS, while for the positive control EGF, a 4.5-fold increase was obtained. When calculating the effect size according to Hedges’ G values, a median and high effect was observed for the EntF* and EGF treatment groups, respectively, compared to the placebo group. [Figure prepared in R-script: https://github.com/JoachimGoedhart/PlotsOfDifferences]

Histopathological data further showed a significant higher number of tumours in both lungs and liver: a significantly higher number of tumour nodules was found in the liver (p=0.014) and lungs (p=0.026) after EntF* treatment for 6 weeks, compared to vehicle treatment (Fig. 5a-d). This is important, as the liver is the most common site of metastases from colorectal cancer: in clinical practice, up to half of all patients with colorectal cancer will develop hepatic metastases, with a median survival of only 8 months and a 5-year survival of less than 5%^26–30^.

**Figure 5:**
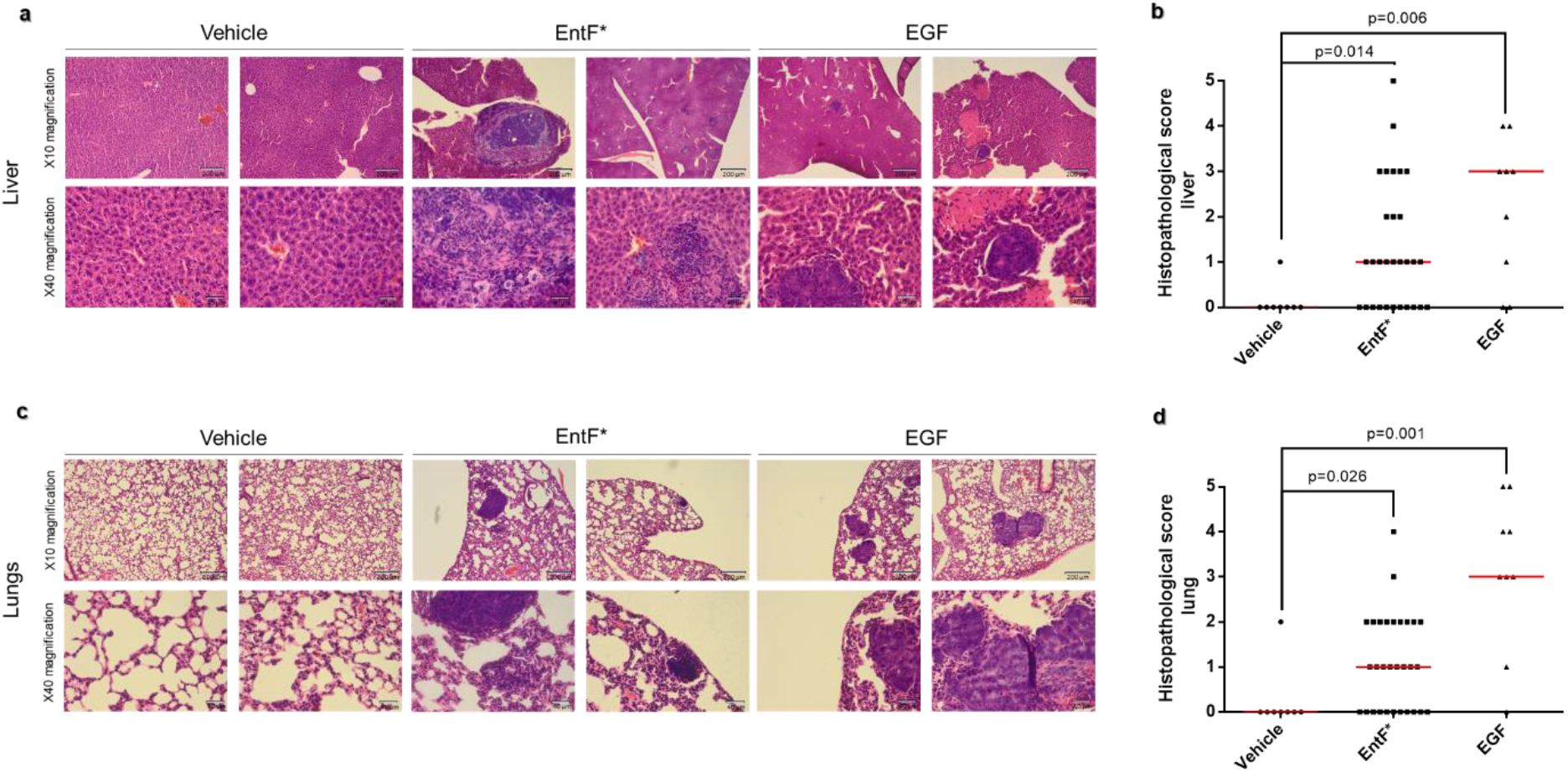
Histopathological evaluation of CRC metastasis after EntF* treatment. **a,** Haematoxylin and eosin (H&E) staining of the liver with magnified images (x10 and x40). **b,** Histopathological scores with statistically significant differences determined by a Mann-Whitney U test (*n*= 8 for PBS, *n*= 30 for EntF*, *n*= 9 for EGF) with indicated p-values. **c,** H&E staining of the lungs with magnified images (x10 and x40). **d,** Histopathological scores with statistically significant differences determined by a Mann-Whitney U test (*n*= 8 for PBS, *n*= 30 for EntF*, *n*= 9 for EGF) with indicated p-values.

Another quorum sensing peptide (*i.e*. Phr0662 from *Bacillus* species), which also promoted *in vitro* cell invasion in our initial screening experiments^13^, showed no *in vivo* metastasis-promoting effects in the same orthotopic colorectal cancer mouse model (Supplementary Fig. 4). These findings indicate the selectivity of the quorum sensing peptides on the *in vivo* metastasis effects.

To evaluate the possible human relevance of our findings, the potential of *E. faecium* bacterial strains from human sources to produce EntF* was investigated. Next to an initial BLAST search, bacterial strains, isolated from human faecal samples, were investigated at DNA level. The results showed that different EntF-containing *E. faecium* strains are present in different human faecal samples (Supplementary Tables 2 and 3), while on the other hand, also *E. faecium* strains without this EntF gene in their genome exist (Supplementary Table 2).

## CONCLUSIONS

Collectively, our findings demonstrate that EntF*, a microbiota-derived quorum sensing peptide metabolite, is present *in vivo* in biofluids of mice and promotes the metastasis of CRC in an orthotopic animal model, with a potency comparable to that of the well-established human colorectal cancer growth factor, EGF. Our findings are the first indication that quorum sensing peptides are an additional factor in microbiota-host interactions potentially influencing CRC metastasis. Our results offer new perspectives in research and development, ultimately offering new possibilities in disease prevention, diagnosis and therapy by selective modulation of the gut microbiome.

## METHODS

### Tissue homogenate preparation

Krebs-Henseleit (KH) buffer (pH 7.4) (Sigma-Aldrich, Belgium) was prepared by dissolving the powdered medium in 900 mL water while stirring. To this solution, 0.3790 g CaCl_2_×2H_2_O and 2.098 g NaHCO_3_ are subsequently added while stirring. NaOH or HCl was used to adjust to pH 7.4. This solution was then further diluted to 1000 mL using ultrapure water.

For the preparation of colon tissue homogenate that was used for the EntF *in vitro* metabolization study, two colons were collected from two C57BL/6 female mice after cervical dislocation. The used mice received no treatment. After cleaning and rinsing the organs using ice-cold KH buffer, the colons were cut in little pieces and transferred into a 15 mL tube to which 5 mL ice-cold KH buffer was added. The colons were then homogenized for 1 minute. After the larger particles were allowed to settle for about 30 minutes at 5°C, approximately 2 mL of the middle layer was dispensed into a 2 mL Eppendorf tube, and stored at −35°C until use. Just before use, the homogenate was diluted to a protein concentration of 0.6 mg/mL.

Faeces homogenate was prepared by collecting faeces from two non-treated C57BL/6 female mice after cervical dislocation. The same procedures as with the colon tissue were performed. Just before use, the homogenate was diluted to a protein concentration of 0.6 mg/mL.

### Peptide adsorption

Due to adsorption of EntF* to different kinds of plastic and glass material, all tubes and containers were coated before use with a BSA-based anti-adsorption solution^24^.

### Metabolization kinetics

500 μL of tissue homogenate and 400 μL of KH buffer were mixed, together with 100 μL of KH buffer (blank) or 1 mg/mL EntF peptide solution (test), all equilibrated and incubated at 37°C. After 0, 5, 10, 30, 60, 120 and 180 minutes, 100 μL aliquots were taken and immediately mixed with 100 μL of 1% V/V trifluoroacetic acid solution in water, heated for 5 min at 95°C, and cooled for 30 min in an ice-bath. After centrifugation at 16,000 g for 30 min at 5°C, supernatants were analysed by LC_1_-MS_1_ for EntF* quantification.

### Cell culture

Caco-2 and luciferase transfected HCT-8/E11 cells were grown in DMEM medium supplied with 10% foetal bovine serum (FBS) and 1% penicillin-streptomycin (10,000 U/mL) solution. The cells were cultured in an incubator set at 37°C and 5% CO2. When confluent, cells were detached using 0.25% trypsin-EDTA.

*E. faecium* strains (LMG 20720, LMG 23236, LMG 15710 and ATCC 8459) were grown overnight at 37°C in BHI medium under aerobic conditions.

### Western Blot analyses

1 x 10^6^ HCT-8 cells were seeded in each well of a 6-well plate. 24 hours post-seeding, cells were treated with EntF* and its synthesised alanine-derived analogues (100 nM) or placebo. For the antagonist study, 1 μM Nef-M1 and EntF*A1 were mixed with EntF* (500 nM). After 24 hours, cells were detached from the surface and lysed with Thermo Scientific RIPA-buffer. The protein concentration was then determined using the modified Lowry protein assay kit, according to the manufacturer’s instructions. All samples were diluted to the same concentration (*i.e*. 4 μg/μL) using water and diluted 1:1 using 2x Laemmli buffer. Next, the samples were boiled for 5 min at 95°C for denaturation, after which centrifugation for 5 min at 16,000g was performed; the supernatant was then used for Western blot analyses. Therefore, proteins (20 μg) were separated using a Bio-Rad Any kD gel (SDS-PAGE) and transferred to a PVDF membrane. Before the membranes were incubated with antibodies, non-specific binding sites were blocked using 5% skimmed milk solution (1 hour). Western blot was performed using an anti-E-cadherin (1/1000) antibody and incubated overnight at 4°C. Signal intensity was normalized against the total protein content in the lanes. Anti-rabbit-HRP antibody was used for detection (1/2000) (1 hour). Finally, the substrate (5 min) was added and the results were analyzed using the Bio-Rad ChemiDoc EZ imager and Image Lab software. TBS buffer with 0.05% Tween 20 was used for washing between the different steps.

### Intestinal permeability

Caco-2 cells were seeded on Transwell polycarbonate membrane filters (0.4 μm pore size) (Corning, Germany) at a density of 2.6 x 10^5^ cells/cm^2^ and the permeability study performed as described by Hubatsch *et al*.^31^. Cells were filled with Hank’s Balanced Salt Solution (HBSS) and the TER values measured before and after the experiment. Peptide solution (1 μM) was added to the apical chamber and 300 μL aliquots taken after 30, 60, 90 and 120 min of incubation. Samples were analysed using LC_1_-MS_1_. Linear curve fitting was used to calculate the apparent permeability coefficient (P_app_).

### Sample collection and preservation

Mice (C57BL/6), possessing their natural microbiome (*i.e*. unmanipulated mice, without any peptide or bacterial administration), were euthanized by cervical dislocation and the blood collected. After standing for 30 min on ice, blood was centrifuged at 1,000 g for 10 min (room temperature). The supernatant (serum) was then transferred and stored at −35°C until use.

After defaecation, two droppings of faeces were immediately collected and put in liquid nitrogen for max. 1 h. The samples were then stored at −80°C until use.

### Sample preparation

50 μL of mice serum was mixed with 150 μL of 0.5% formic acid in acetonitrile. After sonication for 5 min and vortexing for 5 sec, the mixture was heated for 30 sec at 100°C. The solution was again vortexed and centrifuged for 20 min at 20,000 g (4°C). The supernatant was then further purified using solid phase extraction (SPE) on HyperSep C_18_ plates (Thermo Fisher Scientific, Belgium), which were previously conditioned with acetonitrile and equilibrated with 75% acetonitrile in water, containing 0.375% formic acid. After loading 150 μL of the samples, 120 μL eluent was collected and the organic solvents evaporated using nitrogen (1 L/min) for 5 minutes. The resulting solutions were then further diluted with 30 μL of BSA-based anti-adsorption solution, followed by LC-MS analysis.

Bacterial culture medium was centrifuged for 10 min at 2095 g and 4°C, after which the supernatant was filtered through a 0.20 μm filter. For the purification of the culture medium, 200 μL of broth was loaded on an Oasis HLB μElution plate (Waters, Belgium), previously conditioned and equilibrated with acetonitrile and water, respectively. EntF was then eluted from the column using 200 μL of 70% methanol in water containing 2% of formic acid, and the organic solvents evaporated using nitrogen (1 L/min) for 4 minutes. The resulting solution was then further diluted with 150 μL of acetonitrile containing 2% of formic acid, followed by LC_2_-MS_1_ analysis.

### RP-UPLC-TQ-MS (LC_1_-MS_1_) analysis

EntF* was detected and quantified on a Waters Acquity UPLC H-class system, connected to a Waters Xevo™ TQ-S triple quadrupole mass spectrometer with electrospray ionization (operated in positive ionization mode). Autosampler tray and column oven were thermostated at 10°C ± 5°C and 60°C ± 5°C, respectively. Chromatographic separation was achieved on a Waters Acquity^®^ UPLC BEH Peptide C_18_ column (300 Å, 1.7 μm, 2.1 mm x 100 mm). The mobile phases consisted of 93:2:5 water:acetonitrile:DMSO (V/V) containing 0.1% formic acid (*i.e*. mobile phase A) and 2:93:5 water:acetonitrile:DMSO (V/V) containing 0.1% formic acid (*i.e*. mobile phase B), and the flow rate was set to 0.5 mL/min. From the samples, a 10 μL aliquot was injected. The gradient program started with 80% of mobile phase A for 1 minute, followed by a linear gradient to 40% of mobile phase A for 3.5 minutes. Gradient was then changed to 14.2% mobile phase A at 5 min, followed by a 1 min equilibration, before starting conditions were applied. EntF* showed retention at 4.25 – 4.45 min.

An optimised capillary voltage of 3.00 kV, a cone voltage of 20.00 V and a source offset of 50.0 V was used. Acquisition was done in the multiple reaction monitoring (MRM) mode. The selected precursor ion for EntF* was *m/z 865.7* with two selected product ions at *m/z 202.08* (36 eV, b_2_ fragment) as quantifier and *m/z 315.17* (31 eV, b_3_ fragment) as qualifier.

A sample was considered positive for the presence of EntF* when following criteria were met: correct retention time, quantifier/qualifier peak area ratio’s between 2.0 and 4.0, both quantifier and qualifier with a signal-to-noise ratio above 3.0 and a concentration above the LOQ of 100 pM.

### RP-UPLC-QTOF-MS (LC_1_-MS_2_) analysis

Chromatographic separation was achieved on a Waters Acquity^®^ UPLC HSS T3 Column (100 Å, 1.8 μm, 2.1 mm x 100 mm), with detection using the Waters SYNAPT G2-Si High Definition Mass Spectrometry with electrospray ionization (operated in the positive ionization mode). Gradient composition and UPLC-MS settings were the same as with the LC_1_-MS_1_ method; a TOF-MS/MS mode was applied with a fixed mass on the quadrupole of 865.157, a fixed trap collision energy of 30 eV and an acquired MS/MS over the range of 100-1450 m/z (scan time 1 second). EntF* retention was observed between 3.40-3.50 min. When at least four daughter ions (m/z ± 0.05) of EntF* were detected at the expected retention time and at least three of the most abundant isotope parent peaks (m/z ± 0.05) were detected, the sample was considered to contain the EntF* peptide.

### RP-UPLC-QOrbitrap-MS (LC_1_-MS_3_) analysis

While the UPLC separation system was the same as with the LC_1_-MS_1_ method, the third detection system consisted of a Thermo Fisher Q Exactive™ Hybrid Quadrupole-Orbitrap Mass Spectrometer. The mass spectrometer was operated using a heated electrospray ionization source with the following setting: capillary temperature set at 300°C, S-Lens RF level set at 50, spray voltage set at 3.00 kV and auxiliary gas flow set at 20.

A full MS/MS mode was applied with a fixed mass on the quadrupole of 865.157, a fixed trap collision energy of 30 eV and 35 eV and an acquired MS/MS over the range of 100-1800 m/z. EntF* retention was observed at 4.13-4.16 min. When at least two daughter ions (m/z ± 0.005) of EntF* were detected at the expected retention time and at least four of the most abundant isotope parent peaks (m/z ± 0.005) were detected, the sample was considered to be positive for the presence of EntF*.

### HILIC-UPLC-TQ-MS (LC_2_-MS_1_) analysis

Chromatographic separation was achieved on a Waters Acquity^®^ UPLC BEH Amide Column (130 Å, 1.7 μm, 2.1 mm x 100 mm). Mobile phase composition, sample volume, flow rate and MS settings were the same as described for the LC_1_-MS_1_ method. For EntF* quantification in mouse serum, the gradient program started with 10% of mobile phase A for 2 minutes, followed by a linear gradient to 40% of mobile phase A for 3.0 minutes. Gradient was then changed to 85% mobile phase A at 6 min, followed by a 1 min equilibration, before starting conditions were applied. EntF* showed retention at 4.85 – 4.95 min. A sample was considered positive for the presence of EntF* when following criteria were met: correct retention time, both daughter fragments (*i.e*. b_2_ (quantifier) and b_3_ (qualifier) fragment ions) with a signal-to-noise ratio above 3.0 and quantifier/qualifier peak area ratio’s between 2.0 and 4.0.

For the quantification of EntF in culture medium, the gradient program started with 100% of mobile phase B for 2 minutes, followed by a linear gradient to 40% of mobile phase B for 7 minutes, cleaning at 85% B and re-equilibration at starting conditions. Acquisition was done in the multiple reaction monitoring (MRM) mode. The selected precursor ion for EntF was *m/z 667.1* with three selected product ions: *m/z 129.0* (30 eV, b_2_ fragment) and *m/z 662.6* (22 eV, b_2_5 fragment), both as qualifier, and *m/z 949.4* (22 eV, y_17_ fragment) as quantifier.

### DNA extraction of faeces

To 20-40 mg faeces, 500 mg of unwashed glass beads, 0.5 mL CTAB buffer (hexadecyltrimethylammonium bromide 5% (w/v), 0.35 M NaCl, 120 mM K2HPO4) and 0.5 mL phenol-chloroform-isoamyl alcohol mixture (25:24:1) were added. The mixture was homogenized two times for 1.5 min at 22.5 Hz using a TissueLyser II (Qiagen, Belgium). The mixture was centrifuged for 10 minutes at 8,000 rpm and 300 μL of the supernatant was transferred to a new Eppendorf tube. For a second time, 0.25 mL of CTAB buffer was added to the original DNA sample, which was again homogenized in the TissueLyser and centrifuged for 10 minutes at 8,000 rpm. Of this supernatant, 300 μL was added to the first 300 μL supernatant. The phenol was removed by adding an equal volume of chloroform-isoamyl alcohol (24:1) followed by centrifugation at 16,000 g for 10 sec. The aqueous phase was transferred to a new tube. Nucleic acids were precipitated with 2 volumes PEG-6000 solution (polyethyleenglycol 30% (w/v), 1.6 M NaCl) for 2 h at room temperature. The pellet was obtained by centrifugation at 13,000 g for 20 min and washed with 1 mL of ice-cold 70% (v/v) ethanol. After centrifugation at 13,000 g for 20 min, the pellet was dried and resuspended in 50 μL de-ionized water. The quality and the concentration of the DNA was examined spectrophotometrically.

### qPCR on faeces

qPCR was performed using SYBR-green 2x master mix in a Bio-Rad CFX-384 system. Each reaction was done in sixfold in a 12 μL total reaction mixture using 2 μL of the DNA sample and 0.5 μM final qPCR primer concentration. The qPCR conditions used: 1 cycle of 95°C for 10 min, followed by 40 cycles of 95°C for 30 sec, 60°C for 30 sec, and stepwise increase of the temperature from 65° to 95°C (at 10 sec/0.5°C). Melting curve data were analysed to confirm the specificity of the reaction. Samples with aspecific melting peaks were discarded from further analyses. The copy numbers of samples were determined by comparison of their Ct values to the standard curve. For the creation of the standard curves, the PCR product was generated using the standard fragment PCR primers, listed in Supplementary Figure 2, and DNA from *E. faecium* strain 100-1. After purification (MSB Spin PCRapace, Stratec Molecular, Berlin, Germany) and determination of the DNA concentration, the concentration of the linear dsDNA standard was adjusted to 1×10^7^ to 1×10^1^ copies per μL with each step differing by 10-fold. Because the Cq values of the EntF* qPCR analyses were around the limit of detection (LOD), with a notable amount of left-truncated data (data below LOD), a maximum likelihood (ML) approach was used to find the best estimation of mean and standard deviation for each sample.

### Standard protein BLAST

The amino acid sequence of the EntF* peptide was blasted against the NCBI non-redundant (nr) database by Basic Local Alignment Search Tool protein (BLASTp). This blast search was performed with the organism limited to bacteria (taxid:2). Only alignment hits with a 100% coverage and 100% identity were retained.

### Orthotopic colorectal cancer mouse model

All *in vivo* experiments were performed according to the Ethical Committee principles of laboratory animal welfare and approved by our institute (Ghent University, Faculty of Medicine and Health Sciences, approval number ECD 17-90). Mice were maintained in a sterile environment with light, humidity and temperature control (light–dark cycle with light from 7:00 h to 17:00 h, temperature 21–25°C and humidity 45–65%). Before the experiment, mice were allowed to acclimatize for a minimum of seven days.

Six-weeks old female athymic nude mice (Swiss nu/nu) were anesthetized and a small midline laparotomy executed to localize the caecum. The caecum was then gently exteriorized and luciferase transfected HCT-8/E11 cells (1 x 10^6^ cells) in a volume of 20 μL serum-free DMEM medium with matrigel (1:1) injected into the caecal wall. Cells were previously treated with EntF* (10 nM, 100 nM or 1 μM), Phr0662 (100 nM) or with the vehicle (PBS) or positive control (Transforming Growth Factor α (TGFα), 0.1 μg mL^−1^) solution for 5 days before they were implanted in the mice. The caecum was then carefully returned to the abdominal cavity and the laparotomy closed in two layers by sutures of PDS 6/0. Starting from the day after tumour cell injection, mice were daily treated with vehicle (PBS, n = 15), EntF* (10 nmol kg^−1^, n = 12; 100 nmol kg^−1^, n = 18; 1 μmol kg^−1^, n = 8), Phr0662 (100 nmol kg^−1^, n = 5) or positive control (Epidermal Growth Factor (EGF), 100 μg kg^−1^, n = 18) for 6 weeks. Once a week, mice were investigated for tumour growth and metastases using bioluminescent imaging with the IVIS Lumina II (Perkin Elmer, Belgium) after i.p. injection with 200 μL luciferin (150 mg kg^−1^). After 6 weeks, mice were euthanized using cervical dislocation, followed by macroscopic evaluation of the liver, diaphragm, lungs, caecum, duodenum and peritoneum for the presence of tumour nodules. Liver and lung tissues were then fixed in formalin during 24 h and stored in 70% ethanol for max. 3 days before embedding in paraffin. Afterwards, a haematoxylin & eosin (H&E) staining was performed on 8 μm sections and 3 sections were visualized per mouse using microscopy. The slides of all tumour-bearing mice were scored by two blinded, independent investigators using a scoring system as described in Supplementary Table 4. In the case of a difference in scoring, the slide was scored again by a third blinded, independent investigator for consensus.

Daily peptide exposures were calculated after i.p. injection of 25 μL of a 100 μM EntF* solution into female Swiss nu/nu mice (n=14), followed by LC_1_-MS_1_ analyses of mice serum at different time points after injection. A distribution and early exponential phase (α, 0-30 min), followed by a terminal elimination phase (β, 30-180 min) could be distinguished. The exposure was determined for 24 hours (x) as follows: 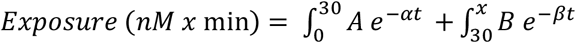.

### PCR on *E. faecium* strains

*E. faecium* was grown in BHI medium. DNA was extracted using alkaline lysis after which the EntF* fragment was amplified using 2x Biomix (Bioline, Belgium) in a Mastercycler PCR system (Eppendorf, Belgium). Each reaction was performed in a 10 μL total reaction mixture using 1 μL of the DNA sample and 0.5 μM final primer concentration (EntF*-PCR primers, Supplementary Fig. 2). The PCR conditions used: 1 cycle of 94°C for 5 min, followed by 30 cycles of 94°C for 30 sec, 55°C for 30 sec and 72°C for 1 min. Final elongation was performed at 72°C for 10 min, after which the PCR product was hold at 4°C. The PCR amplification products were visualized on 1.5% agarose gel.

### Statistical analyses

A quantitative approach was used to evaluate the importance of each amino acid, where the Fisher’s LSD p-values of (1) the multiple comparison between EntF* and the alanine scan and (2) the multiple comparison between different peptides of the alanine scan are combined. Subsequently, based on the combined P-score, the amino acids were classified in 5 classes using a hierarchical cluster analysis, and confirmed using the Jenks natural break algorithm with K=5.

The Kolmogorov-Smirnov test was used to assess if data obtained were normally distributed. For sample sizes of n < 10, non-parametric tests (Mann-Whitney U test) were performed directly. Slope comparison was based on linear regression analysis. Bootstrapped medians and Hedges G-values were used to calculate the effect size when sample sizes were different between the groups. Cohen’s d values were calculated as a measure of the effect size when similar standard deviations for both groups were found and sample sizes were the same.

## DECLARATIONS

### Ethics approval and consent to participate

Not applicable.

### Consent for publication

Not applicable.

### Availability of data and material

Not applicable.

### Author contributions

N.D. and E.W. performed the experiments, with a major contribution on the LC-MS analyses and *in vivo* mice studies. Y.J. and F.V. helped with the Western Blot analyses; A.D.S. and L.T. helped with the qPCR analyses and *in vivo* mice studies, respectively. N.D., Y.J., S.V.W., D.L. and B.D.S designed the Western Blot experiments and discussed the results. N.D., E.W., A.D.S., E.G., F.V.I. and B.D.S. designed the qPCR analyses and evaluated the data. N.D., D.K. and R.H. performed the peptide synthesis of the alanine-derived peptide analogues. E.W., C.V.D.W., O.D.W. and B.D.S. designed the *in vivo* mice studies. N.D., E.W. and B.D.S. wrote the manuscript with input from all co-authors.

## Acknowledgements

We thank the group of Ward De Spiegelaere and Wim Van Den Broeck for helping with the paraffin processing of the tissues.

