## Supplementary information for "The Quorum Sensing Peptide EntF* Promotes Colorectal Cancer Metastasis in Mice: A New Factor in the Microbiome-Host Interaction"

#### 1. Supplementary Figures

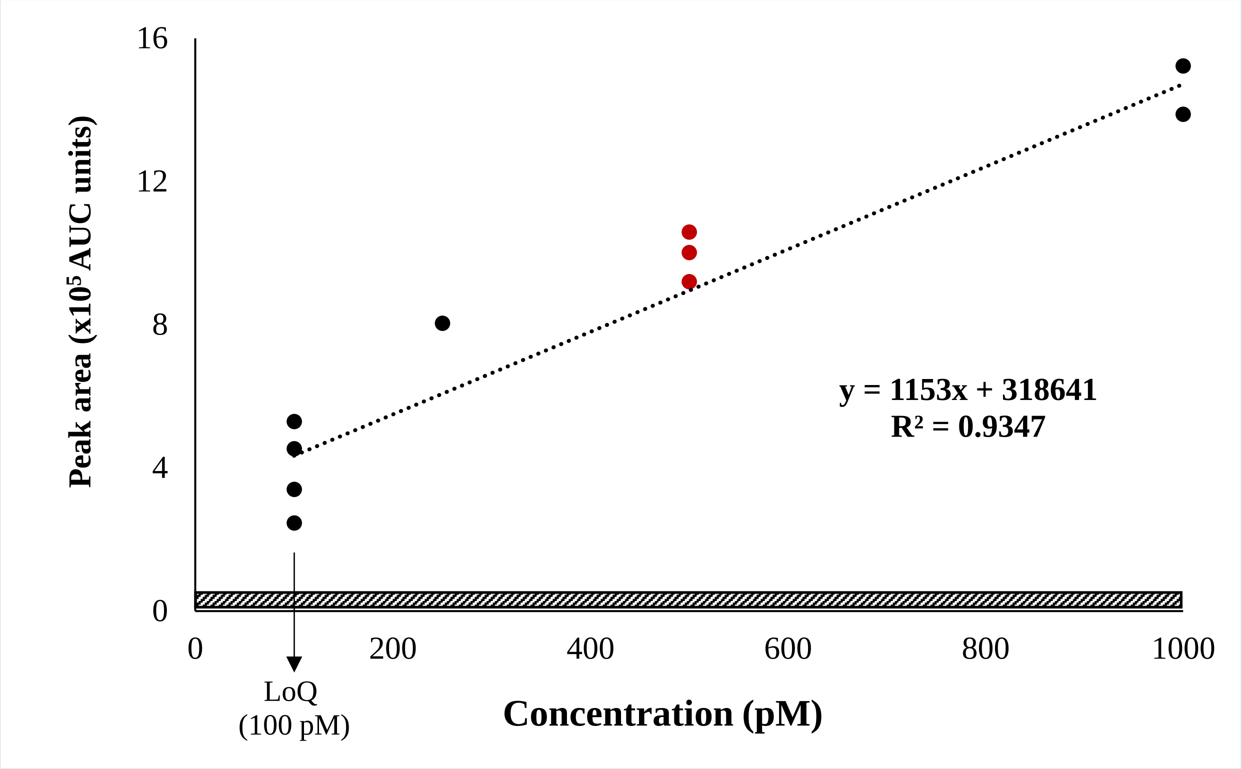


**Supplementary Fig. 1 | Verification of the applied $LC_1$-$MS_1$ method.** The calibration curve was constructed out of 7 independent measurements with independent sample preparation of a serum sample spiked at 3 different concentrations: 100 pM (*n*=4), 250 pM (*n*=1) and 1 nM (*n*=2). The best-fitted regression line represents the calibration curve with indicated $R^2$-value. The accuracy (± 17.1%) and the precision (RSD= ± 10.3%) were determined out of the QC samples measured at 500 pM (*n*=3). A precision of 31.7% was measured at the limit of quantification. The shaded bar represents the area measured in negative samples.

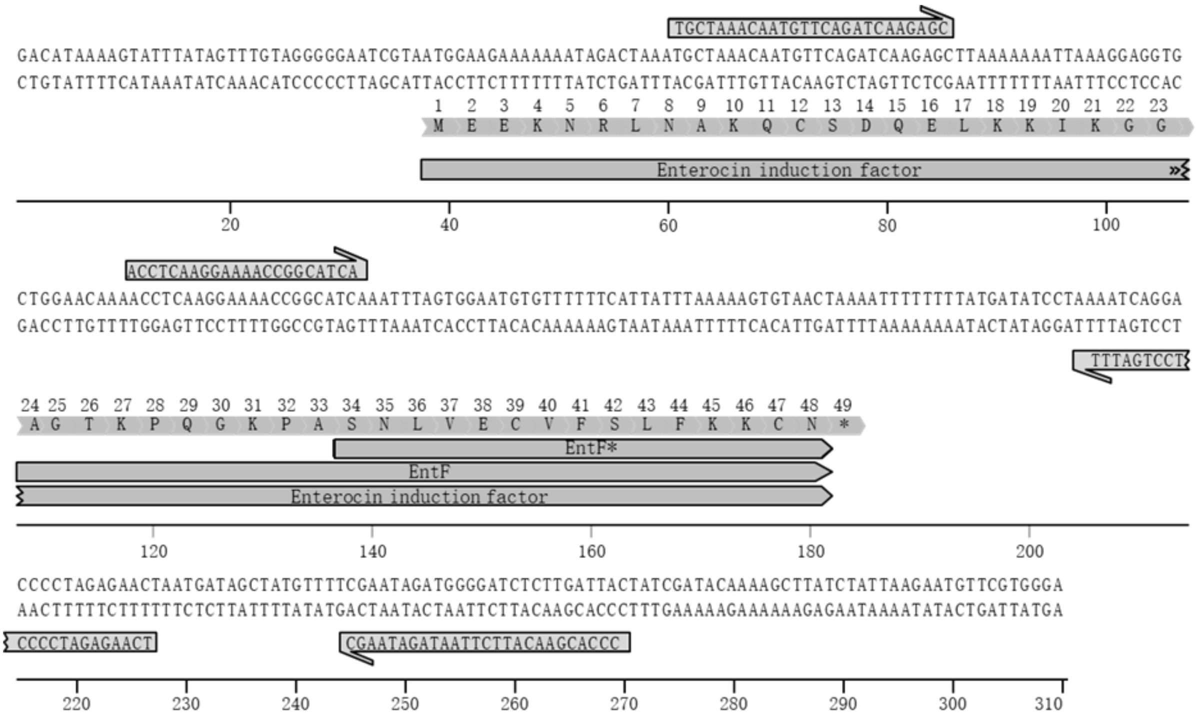


**Supplementary Fig. 2│ qPCR detection of EntF*.** DNA was extracted out of 20-40 mg of faeces and qPCR was performed in sixfold using indicated inner primers. Standard curves were made using the amplicon limited by indicated outer primers.

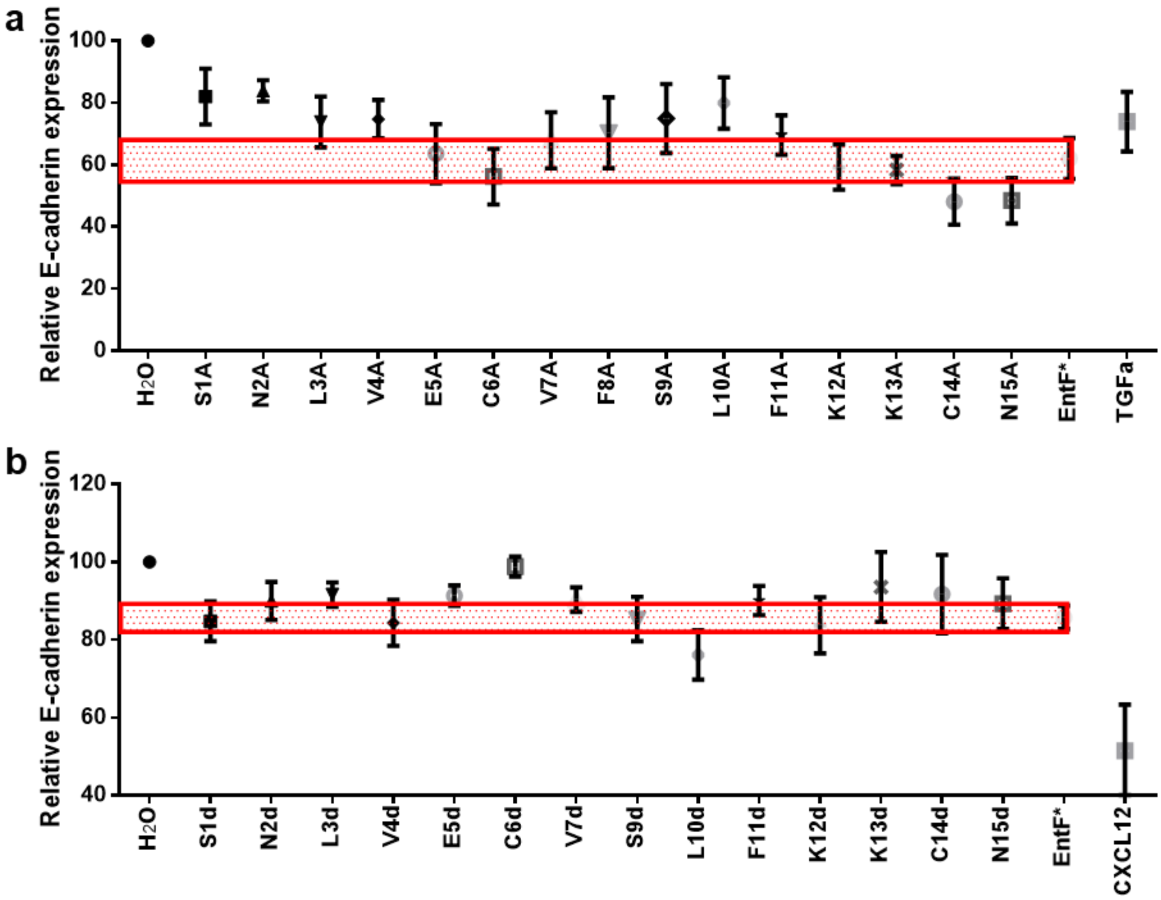


**Supplementary Fig. 3│Effect of EntF* and analogues on E-cadherin expression. a**, Relative E-cadherin levels of ala-scan peptides and EntF* against placebo-treated cells, with TGFa as positive control (mean ± s.e.m.; n=12) during the alanine experiment. **b**, Relative E-cadherin levels of EntF* and its D-amino acid isomers against placebo-treated cells, with CXCL12 as positive control (mean ± s.e.m.; n=11), obtained using Western blotting and densitometric image quantification. Significant different E-cadherin levels were observed between EntF* and EntF*d6, where the sixth amino acid of EntF* was replaced by its D-amino acid isomer.

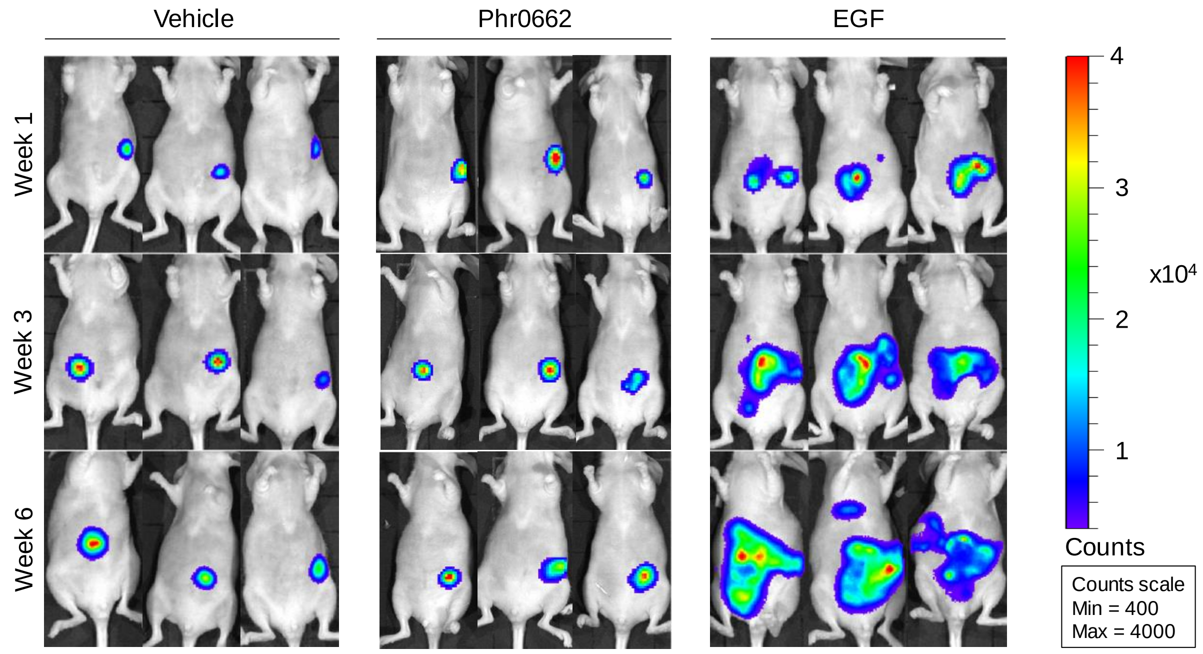


**Supplementary Fig. 4│ The *in vivo* effect of the Phr0662 quorum sensing peptide on colorectal cancer cell metastasis in an orthotopic colorectal cancer mouse model.** A representative image comparing the basal bioluminescence activity between the placebo (n = 7), Phr0662 (100 nmol $kg^{-1}$, n = 5) and EGF (100 µg $kg^{-1}$, n = 8) treatments is given. Mice were i.p. injected with 150 mg $kg^{-1}$ luciferine and imaged 10 minutes later in the supine position. No significant metastasis was observed after daily Phr0662 treatment for 6 weeks, compared to the placebo treatment.

### 1 SUPPLEMENTARY TABLES

| Sample | ID | Detection | Confirmation | | | |
|---|---|---|---|---|---|---|
| | | EntF* concentration (pM) $LC_1$-$MS_1$ | $LC_1$-$MS_2$ | $LC_1$-$MS_3$ | $LC_2$-$MS_1$ | qPCR (copies/g faeces) |
| 1 | 20180827S1 | <LOQ | | | | |
| 2 | 20180827S2 | <LOQ | | | | |
| 3 | 20180827S3 | <LOQ | | | | |
| 4 | 20180827S5 | <LOQ | | | | |
| 5 | 20180827S6 | <LOQ | | | | |
| 6 | 20180827S7 | <LOQ | | | | |
| 7 | 20180827S8 | <LOQ | | | | |
| 8 | 20180827S9 | <LOQ | | | | |
| 9 | 20180827S10 | <LOQ | | | | |
| 10 | 20180829S2 | <LOQ | | | | |
| 11 | 20180829S3 | <LOQ | | | | |
| 12 | 20180829S4 | <LOQ | | | | |
| 13 | 20180829S5 | <LOQ | | | | |
| 14 | 20180829S6 | <LOQ | | | | |
| 15 | 20180829S7 | <LOQ | | | | |
| 16 | 20180829S8 | <LOQ | | | | |
| 17 | 20180829S9 | <LOQ | | | | |
| 18 | 20181011S1 | <LOQ | | | | |
| 19 | 20181011S2 | <LOQ | | | | |
| 20 | 20181011S3 | <LOQ | | | | |
| **21** | **20181011S4** | **389** | | | | |
| 22 | 20181011S5 | <LOQ | | | | |
| 23 | 20181011S6 | <LOQ | | | | |
| **24** | **20181011S7** | **2146** | + | + | + | 588 |
| 25 | 20181011S8 | <LOQ | - | - | - | 612 |
| **26** | **20181011S9** | **3127** | + | + | + | 3060 |
| 27 | 20181011S10 | <LOQ | - | - | - | < LOQ |
| 28 | 20181011S11 | <LOQ | - | - | - | 39 |
| **29** | **20181011S12** | **2489** | + | + | + | 64 |
| 30 | 20181011S13 | <LOQ | - | - | - | 375 |
| 31 | 20181011S14 | <LOQ | | | | |
| 32 | 20181011S15 | <LOQ | | | | |
| **33** | **20181011S16** | **3148** | + | + | + | 1858 |
| 34 | 20181011S17 | <LOQ | | | | |
| **35** | **20181011S18** | **200** | | | | |

**Supplementary Table 1 | Concentration of EntF* measured in mice serum using the $LC_1$-$MS_1$ method and confirmation of the results using additional chromatographic and qPCR**

**methods.** Out of the 35 serum samples, 6 tested positive for the presence of EntF* (indicated in bold), using the $LC_1$-$MS_1$ method. Four positive and four negative samples, indicated in red, were used for the confirmatory experiments. Taking all 35 results into consideration, with a concentration <LOQ set equal to zero, then the average ± s.e.m. is 329 pM ± 150 pM (n=35). If only the six positive samples are considered, then the average ± s.e.m. is 1.91 nM ± 0.54 nM (n=6). The presence or absence of EntF* in the 6 samples was confirmed using the 3 different chromatographic methods. With qPCR, EntF* DNA copies were also observed in all four LC-MS positive samples (*i.e.* 20181011S7, 20181011S9, 20181011S12, 20181011S16); no EntF* copies could be detected in sample 20181011S10. +: present; -: not present; grey: not investigated.

| **Bacterial strain** | **Origin** | **EntF gene present?** | **EntF peptide present?** |
|---|---|---|---|
| *E. faecium* LMG 20720 | Human faeces | Yes | < LOD |
| *E. faecium* LMG 23236 | Human faeces (healthy) | Yes | < LOD |
| *E. faecium* LMG 15710 | Human faeces (diarrhea) | No | < LOD |
| *E. faecium* ATCC 8459 | Dairy product (cheese) | Yes | 477 nM |

**Supplementary Table 2│Presence of EntF gene and peptide in different *E. faecium* strains.** Four different, commercially available strains of *E. faecium* were investigated for the presence of the EntF gene using PCR. From three out of the four strains, the EntF gene was confirmed, while this was absent for the LMG 15710 culture. The presence of the EntF peptide in the culture media was also investigated: out of the three *E. faecium* strains that contain the EntF gene, only one strain produced EntF *in vitro* (LoD = 1.5 nM).

| EntF producing strains found in human faeces | |
|---|---|
| *E. faecium* LIM714,<br>*E. faecium* VRE-1402237,<br>*E. faecium* U-1313438,<br>*E. faecium* E1071,<br>*E. faecium* ERV165,<br>*E. faecium* GMD5E,<br>*E. faecium* GMD4E,<br>*E. faecium* GMD3E,<br>*E. faecium* GMD2E,<br>*E. faecium* GMD1E,<br>*E. faecium* 6E6,<br>*E. faecium* MXVK29,<br>*E. faecium* S-1001508,<br>*E. faecium* VRE-1402513,<br>*E. faecium* VRE-1402563,<br>*E. faecium* VRE-1406033,<br>*E. faecium* VRE-1504220,<br>*E. faecium* LMG 8148,<br>*E. faecium* Isolate 30,<br>*E. faecium* VRE16,<br>*E. faecium* XH877,<br>*E. faecium* HMSC063H10,<br>*E. faecium* HMSC069A01,<br>*E. faecium* HMSC056C08,<br>*E. faecium* 97-7_S6,<br>*E. faecium* LIM1546,<br>*E. faecium* LIM1547,<br>*E. faecium* LIM1759, | *E. faecium* 2014-VREF-41,<br>*E. faecium* 2014-VREF-63,<br>*E. faecium* 2014-VREF-26,<br>*E. faecium* F1129F 09,<br>*E. faecium* F1213D 01,<br>*E. faecium* 24,<br>*E. faecium* Hp_24-3,<br>*E. faecium* Hp_5-10,<br>*E. faecium* Hp_6-10,<br>*E. faecium* Hp_5-7,<br>*E. faecium* 8_Efcm_HA-DE,<br>*E. faecium* 11_Efcm_HA-DE,<br>*E. faecium* FDAARGOS_326,<br>*E. faecium* 4278,<br>*E. faecium* 1584,<br>*E. faecium* IHC105,<br>*E. faecium* IHC116,<br>*E. faecium* IHC106,<br>*E. faecium* IHC113,<br>*E. faecium* IHC115,<br>*E. faecium* IHC130,<br>*E. faecium* IHC117,<br>*E. faecium* IHC121,<br>*E. faecium* IHC118,<br>*E. faecium* IHC127,<br>*E. faecium* IHC108,<br>*E. faecium* IHC120,<br>*E. faecium* IHC123 |

**Supplementary Table 3 │ Bacterial strains found in human faeces samples with indicated presence of the EntF gene in the genome.** Using BLASTp searches, the isolation source was investigated for the resulting *E. faecium* strains, with different strains found in human faeces samples.

| Liver scoring system | | | |
|---|---|---|---|
| **Score** | **Signs** | **Representation x10** | **Representation x40** |
| 0 | • Well-structured cells<br>• Absence of nodular infiltrates<br>• Absence of necrotic tissue | 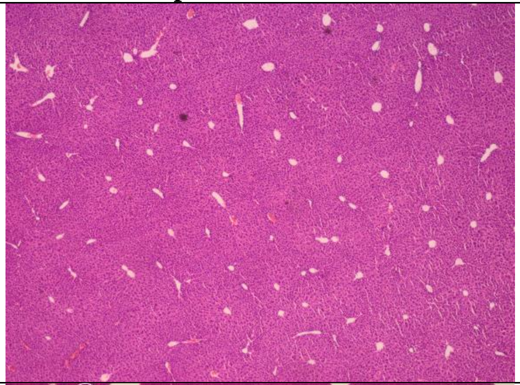 | 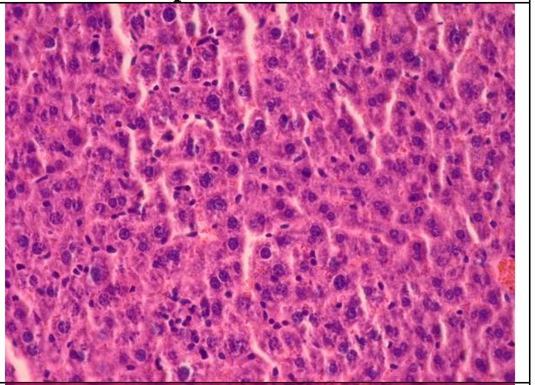 |
| 1 | • Nodular infiltrates<br>• No capsular organisation<br>• Absence of necrotic tissue | 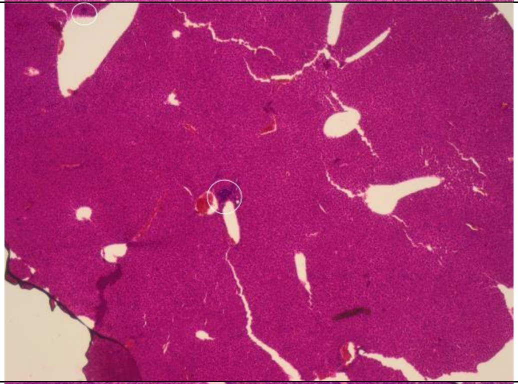 | 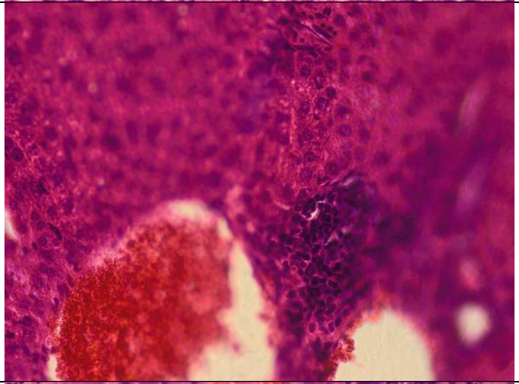 |
| 2 | • Nodular infiltrates<br>• Capsular organisation around infiltrates<br>• Absence necrotic tissue | 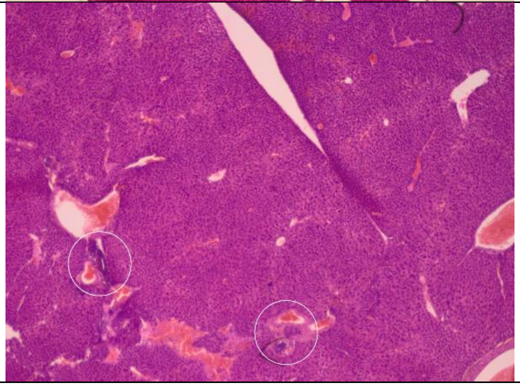 | 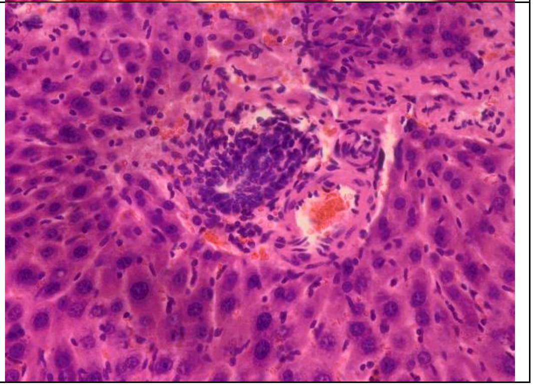 |

| | | | |
|---|---|---|---|
| 3 | • Nodular infiltrates<br>• Presence of necrotic tissue | 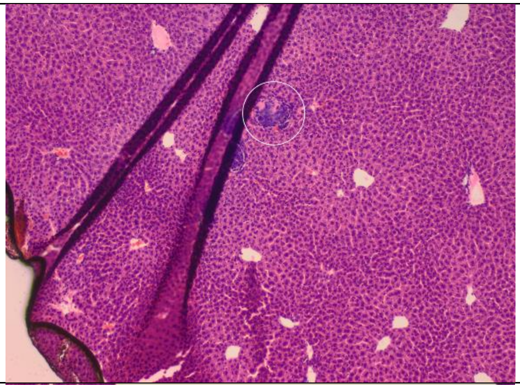 | 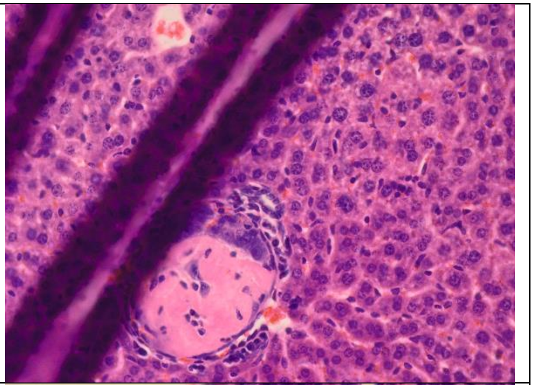 |
| 4 | • Large nodular infiltrates<br>• Large patches of necrotic tissue in nodules<br>• Clear distinct capsular organisation<br>• Less than ¼ of the liver coupe consist of tumour | 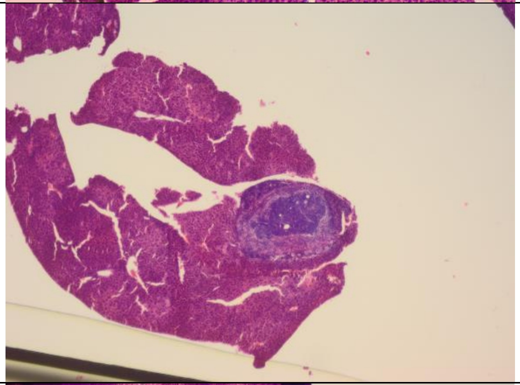 | 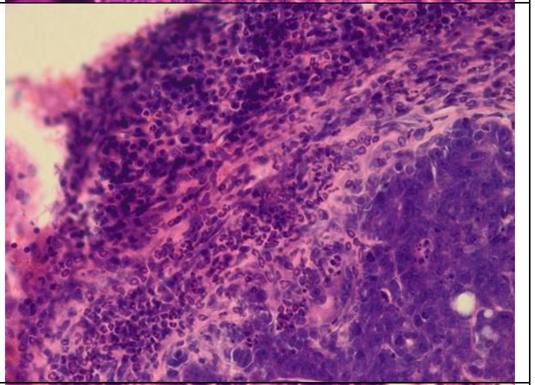 |
| 5 | • Large nodular infiltrates<br>• Large patches of necrotic tissue in nodules<br>• Clear distinct capsular organisation<br>• More than ¼ of the liver coupe consist of tumour | 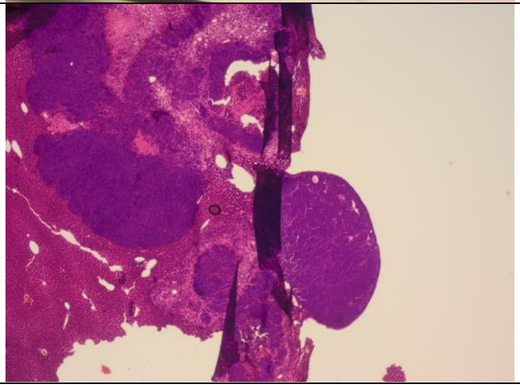 | 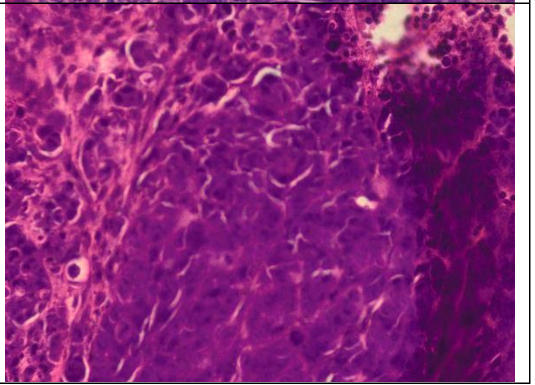 |

| Lung scoring system | | | |
|---|---|---|---|
| **Score** | **Signs** | **Representation x10** | **Representation x40** |
| 0 | • Well-structured cells<br>• Absence of tumour nodule(s)<br>• Absence of necrotic tissue | 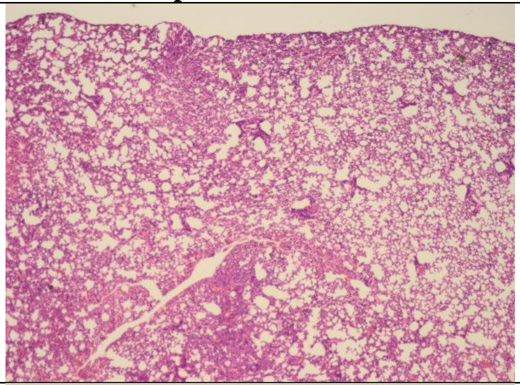 | |
| 1 | • Nodular loose infiltrates<br>• Absence of necrotic tissue | 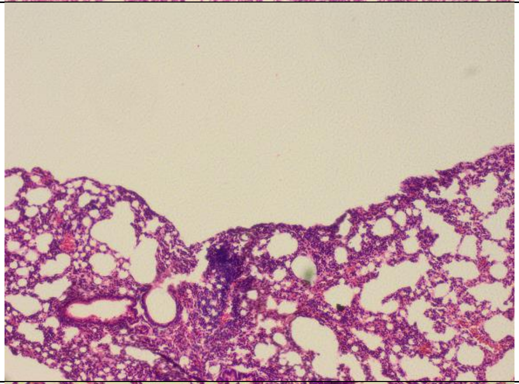 | 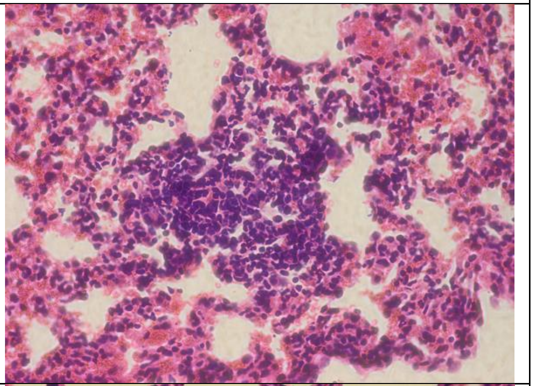 |
| 2 | • Nodular dens infiltrates<br>• Decrease of cytoplasm/nucleus ratio<br>• Absence necrotic tissue | 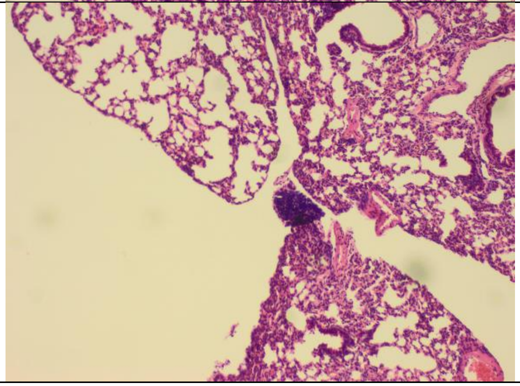 | 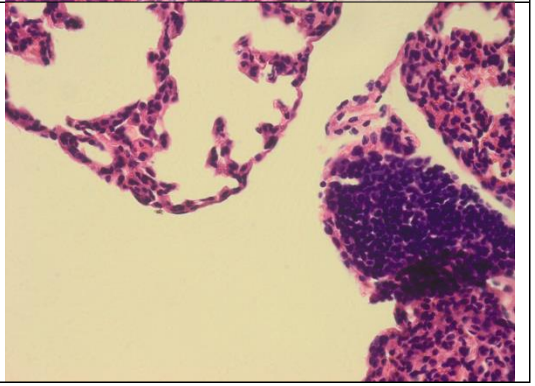 |

| | | | |
|---|---|---|---|
| 3 | • Large nodular infiltrates<br>• Cytoplasmic basophilia<br>• Absence necrotic tissue | 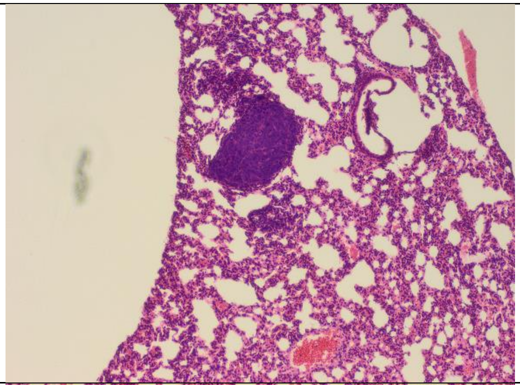 | 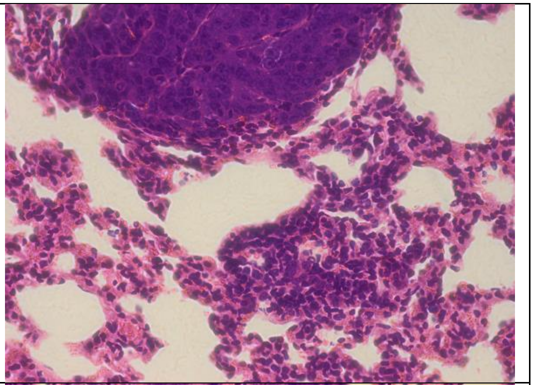 |
| 4 | • Nodular infiltrates<br>• Presence necrotic tissue | 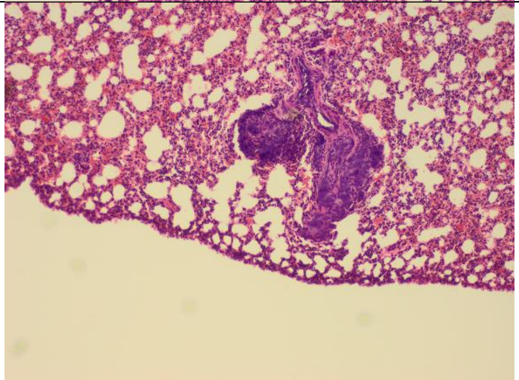 | 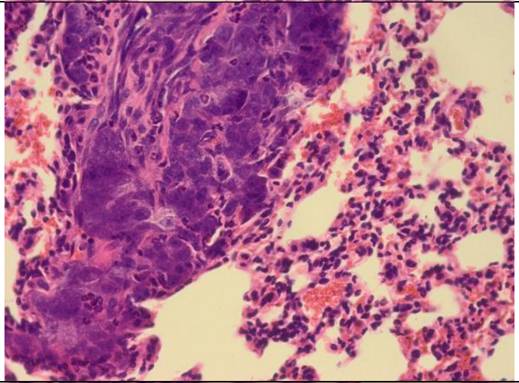 |
| 5 | • > 1 necrotic infiltrate | 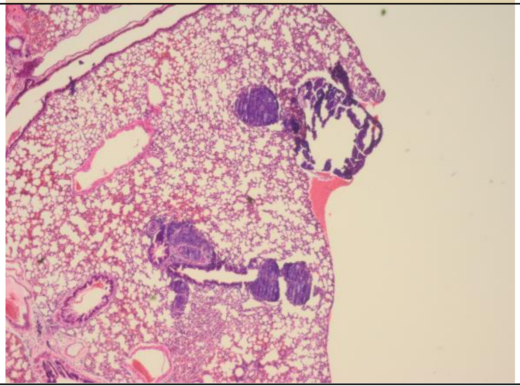 | 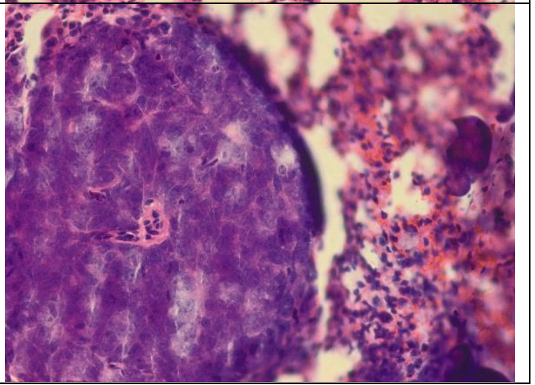 |

**Supplementary Table 4│ Histopathological scoring system.** For both the liver and lung tissue, a scoring system was developed, thereby evaluating the presence and severity of tumour nodules and necrotic tissue. Normal tissue was given a score = 0, while the presence of clear (large) and numerous tumour nodules and necrotic tissues was given a score = 5.
